# The effect of alien species on plant-pollinator network structure across a gradient of plant species invasion

**DOI:** 10.1101/655522

**Authors:** Víctor Parra-Tabla, Diego Angulo-Pérez, Cristopher Albor, María José Campos-Navarrete, Juan Tun-Garrido, Paula Sosenski, Conchita Alonso, Tia-Lynn Ashman, Gerardo Arceo-Gómez

**Affiliations:** Departamento de Ecología Tropical, Campus de Ciencias Biológicas y Agropecuarias, Universidad Autónoma de Yucatán, 97000 Mérida, Yucatán, México; División de Estudios de Posgrado e Investigación, Instituto Tecnológico de Tizimín, Tecnológico Nacional de México, 97700 Tizimín, Yucatán, México; Departamento de Botánica, Campus de Ciencias Biológicas y Agropecuarias, Universidad Autónoma de Yucatán, Km. 15.5 Carretera Mérida-Xtmakuil, Mérida, Yucatán, México, 97000; Estación Biológica de Doñana, Consejo Superior de Investigaciones Científicas (CSIC), Avda. Américo Vespucio s/n, E-41092, Sevilla, Spain; Department of Biological Sciences, University of Pittsburgh, Pittsburgh, PA 15260, USA; Department of Biological Sciences, East Tennessee State University, Johnson City, TN 37614, USA

**Keywords:** coastal communities, conservation, alien species, plant-pollinator networks, pollinator sharing, pollinator competition, sand dunes

## Abstract

The interactions between pairs of native and alien plants via shared use of pollinators have been widely studied. Studies of invasive species effects at the community level on the other hand are still scarce. Few community level studies, however, have considered how differences in the intensity of invasion, and degree of floral trait similarity between native and invasive species, can mediated effects on native plant-pollinator communities. Here, we evaluated the effect of alien species on overall plant-pollinator network structure, and species-level network parameters, across nine coastal communities distributed along 205 km at Yucatán, México that vary in alien species richness and flower abundance. We further assessed the effect of alien plant species on plant-pollinator network structure and robustness via computational simulation of native and invasive plant extinction scenarios. We did not find significant differences between native and alien species in functional floral phenotypes, the visitation rate and species composition of the pollinator community. Variation in the proportion of alien plant species and flower abundance across sites did not affect plant-pollinator networks structure. Species-level network parameters (i.e., normalized degree and nestedness contribution) did not differ between native and alien species. Furthermore, our simulation analyses revealed that alien species are functionally equivalent to native species and contribute equally to network structure and robustness. Overall, our results suggest that alien species are well integrated into native coastal plant-pollinator networks which may be facilitated by high levels of floral trait similarity and pollinator use overlap. As a result, alien species may play a similar role than that of natives in the structure and stability of native plant and pollinator communities in the studied coastal sand dune ecosystem.

## Introduction

Alien plant species can alter a vital ecosystem function by disrupting mutualistic interactions between native plant species and their pollinating partners [1-3]. Alien plants can decrease floral visitation, pollen deposition and reproductive success of native plant species [4-6; but see 7] and alter the structure of community-level plant-pollinator interactions (i.e. network structure; [8-11]. These effects can in turn affect the long-term stability and functionality of native plant communities [12-14]. However, current understanding of alien species effects still strongly relies on studies of interactions between the invasive and one, or very few, native species [4, 7], typically at a single location [8, but see 15-17]. Knowledge on the effect of alien species on the structure and function of entire plant and pollinator communities, and how these effects vary spatially, is needed if we aim to fully understand the effects and consequences of plant invasions in nature [18].

The few community-level studies conducted to date have shown that alien species can alter plant-pollinator network structure. These effects however can vary spatially depending on the pollinator assemblage and the degree of plant and/or pollinator specialization [17]. Less studied however, is how invasive species effects may vary with varying intensity of plant invasion (e.g., alien flower abundance), even though it is unlikely that all communities are equally invaded and that highly invaded communities will respond similarly to low invaded ones [18]. For instance, it has been shown that network connectance and the evenness of interactions within a network is strongly diminished in areas where the proportion of alien flowers are higher than 50% [16], thus suggesting potential density-dependent effects of alien species on native plant-pollinator communities. Furthermore, studies on the impacts of alien species on plant-pollinator network structure have so far shown contrasting results. While some have shown that alien plant species can affect network specialization, modularity (i.e. tight subsets of interacting species; [11, 16]), nestedness and robustness [19, 20], others have found little to no effect on network structure [5, 21]. This apparent discrepancy in the effects of invasive species on network structure may stem from differences in the amount of invasive floral resources available (i.e. intensity of invasion) among study systems, but this has been seldom considered [16, 18]. In order to gain a more complete understanding of the consequences of alien species effects on native plant-pollinator interactions it is thus imperative to evaluate how their effects vary across various levels of alien species richness and floral abundance [17, 22]. Such knowledge may also help explain the conditions that allow alien species to rapidly integrate into native pollination networks [5, 18]. It is also important to note, that even in the absence of alien species effects on overall network structure, alien plants can reduce pollinator visitation and reproductive success of individual community members, negatively impacting native plant communities [11, 17]. Nonetheless, studies that simultaneously consider both, network structure and species-level effects are scarce, thus limiting our understanding of overall alien species effects on native plant-pollinator communities.

The effects of alien species on the pollination of native plants has been often attributed to their generalized pollination system which increases the likelihood of pollinator sharing with natives [5, 11, 18, but see 23]. Floral trait similarity between natives and alien species on the other hand, can further increase the degree of pollinator sharing and augment alien species effects on the pollination success of native plants [5, 21, 24]. For instance, some authors have found that floral trait similarity explained the amount of flower visitor overlap between native and invasive species [24]. Such overlap in pollinator use has the potential to alter pollinator preference [25, 26] and modify the richness and abundance of floral visitors visiting native species [27-29], ultimately affecting the structure of native plant-pollinator networks [12, 17, 22]. While most studies evaluating the importance of floral similarity in mediating interactions between native and alien species have focused on comparisons between species pairs [15, 17, 24], the degree and importance of floral trait similarity between native and alien plants at the community-level has received considerably less attention. Such studies are crucial if we aim to fully understand the factors and mechanisms that facilitate successful plant invasion.

In this study we analyzed the structure of plant-pollinator networks in nine coastal sand dune plant communities distributed along 205 Km of coast in the Yucatan Peninsula, Mexico. These coastal areas, as the vast majority of them worldwide, have been subject to intense human use (e.g. tourism), which has increased the presence of invasive species [30]. Specifically, in the last 30 years, the number of alien species present in the north coast of Yucatan has grown substantially [31, 32]. Currently, almost 30% (20 species) of total plant species richness in this coastal ecosystem is composed of alien species [32]. However, their distribution is not homogeneous along the coast, and some areas remain significantly more invaded compared to others. Site differences in the proportion of alien plant species present range between 22% and 50% [32], while alien floral abundance can range from 11% to 99% across sites (data from this study). Furthermore, seventeen of the alien species present in this ecosystem have been described as pollination generalists [33], and thus their potential for altering native plant-pollinator interaction networks is high [3, 34]. In this study, we characterize and compare the ‘pollination environment’ (i.e. pollinator richness and composition and floral visitation rate) of natives and alien plants. We further characterize plant-pollinator interaction network structure in these nine coastal co-flowering communities to evaluate the effect of alien species proportional richness and floral abundance on network structure. We also evaluate the degree of floral trait similarity between native and alien species across all nine co-flowering communities. Finally, we further assessed the effect of plant species invasion on plant-pollinator network structure and robustness via computational simulations of native and alien plant extinction scenarios. We specifically ask the following questions: 1) Do native and alien plant species differ in pollinator visitation rate and in the diversity and species composition of flower visiting insects, and are differences consistent across the sites? 2) What is the effect of increasing proportion of alien species richness or alien flower abundance on plant-pollinator network structure? 3) What is the degree of floral trait similarity between invasive and alien species in all nine studied communities? and 4) Does simulated extinction of invasive species have the same effect on plant-pollinator network structure compared to extinction of natives or random extinction of species?

## Methods

### Study sites

We studied nine sand dune plant communities distributed along 205 km of coast in northern Yucatan (S1 Fig). The sand dune ecosystem is continuous along the entire coast, which extends over approximately 320km but is interrupted in a few areas by mangrove and lagoon systems [35]. Thus, the studied area encompasses the full distribution of the sand dune ecosystem in the north coast of the Peninsula. We selected nine sites (i.e. co-flowering communities) with different “levels” of invasiveness previously identified along the coast ([32], Table1). Average proportion of alien species (number of alien species /total number of species) across sites ranged between 22% and 50% (average 34.4% ± 10.6; [32]). Proportional alien flower abundance ranged between 11% and 99% (Table 1). Average distance between sites is 19.3 km with a minimum of 4.7 km (S1 Fig).

**Table 1.** Names of study sites. The number of native and alien species (percentage), proportion of alien flowers (percentage) and floral traits similarity between native and alien plant species at each site is shown. Sites are ordered according to proportion of alien flower abundance

| Site | Native species | Alien species (%) | Proportion of alien flowers (%) | Floral traits similarity (mean ± SD) |
| --- | --- | --- | --- | --- |
| Chapo 1 | 9 | 3 (25) | 11 | $0.80 \pm 0.04$ |
| Playa Maya | 7 | 2 (22) | 40 | $0.79 \pm 0.04$ |
| Chapo 2 | 10 | 4 (28) | 78 | $0.77 \pm 0.11$ |
| Telchac | 8 | 6 (42) | 79 | $0.78 \pm 0.05$ |
| Cancunito | 4 | 4 (50) | 90 | $0.80 \pm 0.03$ |
| Punta Meco | 8 | 6 (42) | 92 | $0.79 \pm 0.03$ |
| Sisal | 9 | 6 (27) | 96 | $0.78 \pm 0.04$ |
| Charcas | 9 | 3 (27) | 97 | $0.81 \pm 0.03$ |
| Chabihau | 11 | 9 (47) | 99 | $0.79 \pm 0.04$ |

### Pollination environment and floral traits

We characterize plant and pollinator communities at each site. For this, we established ten 20 m2 plots at each site, each one separated by 20 m. Within each plot we recorded the number and identity of all flowering species and the total number of open flowers per species. In species with numerous small flowers (<1 cm; i.e. Amaranthacea) we estimated the number of flowers as: average number of flowers on three inflorescences × total number of inflorescences in a plant. This estimation was done for eight species (*Atriplex tampicensis, Alternanthera microcephala, Amaranthus greggii, Suaeda linearis, Bidens pilosa, Flaveria linearis* and *Melanthera nivea)*. For each flowering species we also measured the following floral traits: flower height (distance between the calyx and the tip of the corolla), corolla diameter (larger perpendicular distance to height), and corolla opening (opening of the corolla tube; coded as zero in non-tubular species). We measured flower traits in 1-5 flowers per plant in at least three plants per species. We selected these floral characteristics because they have been described as strong mediators of plant-pollinator interactions. For instance, flower height and corolla diameter are traits related to long-distance perception of flowers by pollinators [36, 37] whereas corolla opening is associated with degree of flower generalization (e.g., flowers with small corolla opening have restricted access to floral rewards than flower with larger ones; [38]. We also measured floral reflectance spectra (300-700 nm) of the dominant corolla color in 1-3 flowers per species. Floral reflectance was measured with a field spectrophotometer (StellarNet INC). With this data we estimated flower color using chromatic coordinates (X and Y) of the Hymenoptera vision model [39], which are the most abundant floral visitors in these communities [40, 41]. Estimation of the color-hexagon vision model were carried out with the *pavo* package in the R 3.2 software [42]. With these flower trait data, we estimated floral trait similarity between native and alien plant species in each co-flowering community using Gower’s pairwise distances [24, 37, 41]. We calculated a similarity index for each species (1 – average similarity Gower’s pairwise distances) that represents the degree of similarity of each species (alien or native) with respect to all the species present in the community [24]. An average floral trait similarity index between native and invasive species was estimated for all nine communities.

### Plant-pollinators networks

We characterized plant-pollinator interactions at each site by observing each plot for five minutes, and recording the number and identity (species/morphospecies) of floral visitors for every species flowering within the plot. Only visits in which the insects contacted the reproductive structures of the flowers were recorded. We collected 5-10 specimens of each pollinator species or morphospecies using an entomological net and stored it in microcentrifuge tubes (with the exception of Lepidoptera) for subsequent identification in the laboratory. The short height of the vegetation (< 50 cm tall) and the low density of plants allowed us to accurately observe and record all plant-pollinator interactions within each plot. Each plot was observed three times per day for a total of 15 min plot/day and a total of 150 min per site/day. All observations were carried out during peak pollinator activity between the 0800 and 1200 [40]. Each site was visited a total of nine days (approx. every 10 days). The order in which sites were visited was randomized. Observations were always conducted by the same group of people and insect identification was cross-checked. The study was carried out during peak flowering time in these communities (September to November) in 2016.

## Statistical analyses

### Pollination environment and floral traits

To evaluate differences in pollinator species composition between native and alien plant species, and among sites, we performed a Permutational Multivarate Analysis of Variance (PERMANOVA, [43]). For this, we performed 1000 random permutations based on a distance matrix using pollinator species abundance data calculated with the Bray-Curtis index, and considering the dates as subsamples of each site. These analyses were performed with the Vegan package of R v.3.0.2 [44].

To evaluate differences in average pollinator species richness and pollinator visitation rate between native and alien plant species and among sites (i.e. along the invasion gradient) a mixed GLMM model was performed using site and species origin (i.e., native *vs.* alien) nested in site as fixed effects, and species as a random effect. In these models we included floral abundance (log transformed) as a covariate and used a Poisson and lognormal error distribution for pollinator richness and pollinator visit rate respectively. These analyses were carried out using the GLIMMIX procedure in SAS [45].

To evaluate the degree of floral trait similarity between native and invasive plants we evaluated if alien species were more similar to the pool of all species in the community than the native species [24]. We conducted an independent GLM to test for overall differences in floral trait similarity between alien and native species across all communities and to test for differences within each community. These tests were carried out using the GLM procedure in SAS [45].

### Plant-pollinators networks

To characterize plant-pollinator network structure at each site we constructed an interaction frequency matrix for each site using the number of times every floral visitor was observed visiting flowers of a particular plant species [46]. We then used these matrixes to construct a plant–pollinator network and to estimate the network’s metrics for each site using the ‘bipartite’package in R [43, 47]. We focus on the following specific descriptors of overall network structure: (a) nestedness (non-random interaction pattern among species where specialists interact with subsets of species interacting with generalists; [46]), (b) network-level specialization (H2 index; takes values from 0 [no specialization] to 1 [total specialization] and is suitable for comparisons across different networks; [48], (c) modularity (indicates whether species are structured into subsets that are more strongly connected to one another than to species outside the module), and (d) robustness (estimates the robustness of the system to species loss; [19,20]. Robustness’ values close to 1 indicate networks will be less affected by extinction events [20].

For each network, we evaluated if nestedness differed significantly from random using the NODF parameter (1000 simulations) in Aninhado [49]. We estimated network modularity using the M index which ranges between 0 (no modularity) and 1 (complete modularity) using Simulated Annealing in MODULAR [50, 51]. Random matrices were generated to test the significance of modularity according to the null Model III (CE) using 100 randomizations per network [51]. In this null model the probability of occurrence of an interaction is proportional to the number of interactions of plants and pollinators [52].

Sampling completeness of each plant-floral visitor network (i.e., observed numbers of pairwise plant-floral visitor associations) was verified via rarefaction analysis using EstimateS 9.1 [53]. Rarefaction curves were constructed with 500 randomizations and sampling without replacement [53]. We further calculated an estimator of asymptotic interaction richness (Chao 1) and estimated the percentage of interaction richness detected in our sampling by dividing the observed by the estimated number of pairwise interactions. Average sampling completeness across sites was of 70.2% (± 10.2, SD), suggesting that our sampling sufficiently captured the majority of the expected plant-pollinator interactions. To evaluate if observed network structural parameters vary with degree of invasion, we regressed each network parameter separately on the proportion of invasive plant species, and the proportion (log-transformed) of invasive flowers.

### Species level analysis

For each networks the following species level metrics were calculated: (a) interaction strength (sum of individual dependencies for each species), (b) normalized degree (sum of links per species relative to the total number of possible interacting partners) and (c) nestedness contribution (estimates the individual contribution of each species to overall nestedness; [54]. Strength and normalized degree were calculated using the species level function in the bipartite R package [47], and nestedness contribution [55]. In order to evaluate differences in species level metrics among sites and between native and alien plant species a GLMM was conducted considering site and plant origin (nested in site) as fixed effects, and species as a random effect. Log transformed flower abundance was considered as a covariate. These analyses were carried out using the GLIMMIX procedure in SAS [45].

### Effect of alien species on plant-pollinator network structure via simulation of extinction scenarios

To further evaluate the role of alien plant species on network topology and species-level network estimators three plant species extinction scenarios were simulated: (a) An “aliens removed” scenario, in which all alien plant species present at each site were excluded from the interaction matrix, (b) A “natives removed” scenario, in which we randomly excluded native plant species, and (c) A “random removal” scenario, in which plant species were excluded randomly without consideration about their origin (i.e., native or alien). For the latter two scenarios we removed as many plant species from the network as the number alien species at each site. We compared the structure of these networks with that of (d) “intact” (observed) networks that included all plant species at a site (i.e., natives + aliens). By comparing the structure of “intact” networks to those were only aliens were removed (i.e. “aliens removed”) we evaluated the potential effect of alien species on network structure [19]. Furthermore, by comparing network structure between “aliens removed” vs. “native removed” and “random removal” extinction scenarios, we evaluated if the effect of removing alien species was equivalent to that of removing only native species or from a completely random extinction scenario. Because species exclusion from a matrix modifies both, the number of interactions and connections within the network we used the second.extinct function in R [19, 47] which considers interaction rewiring within the network (i.e., ability of pollinators to visit other plant species when its preferred species is absent; see [11, 16, 40]). Even though the species extinction scenarios simulated here may differ from those in nature, (e.g. because not all plant species may require animal pollination to persist) they allowed us to assess the potential effects of alien species in network structure and their equivalence with natives [11, 19].

To statistically evaluate the differences in network structure among the ‘extinction scenarios’, we conducted mixed GLMM models with extinction scenario (‘intact’, ‘aliens removed’, ‘natives removed’ and ‘random’) as a fixed effect and site as random effect. We evaluated differences on the following network-level metrics: network specialization (H2), modularity, nestedness, and robustness. For network specialization (H2), nestedness, and modularity log normal errors were used. A normal error distribution was used for robustness. Analyses were carried out in SAS [45]. Subsequently, in order to compare the magnitude of change (Δ) in network structure due to species removal in each extinction scenario relative to the ‘intact network’ (observed network), we calculated the bias corrected Hedges’ g effect size: Hedges’g= *M*_1_-*M*_2_/*SD* _pooled_. Where *M*_1_= mean of each network metric observed across communities and *M*_2_= mean of each network metric calculated for each simulated extinction scenario across communities, and *SD* _pooled_= pooled and weighted standard deviations [56]. We performed this for network specialization (H2), modularity, nestedness, and robustness.

To evaluate differences on individual species’ roles (i.e. species-level parameters) within the network among ‘extinction scenarios’, we conducted GLMMs with extinction scenario and plant origin (nested in site) as fixed effects, and species as a random effect on the following species-level parameters: strength, normalized degree and nestedness contribution. A lognormal error was used for normalized degree and nestedness contribution and a Poisson error was used for strength. These analyses were carried out using the GLIMMIX procedure in SAS [45]. In all cases mean ± SD are presented, unless otherwise specified.

## Results

### Pollination environment and floral trait similarity between native and alien plant species

A total of 516609 flowers were recorded. Among-site variation in the total number of open flowers ranged from 31917 (site Chapo 2) to 218206 (site Chabihau) (Fig 1) across the season.

**Fig. 1.**
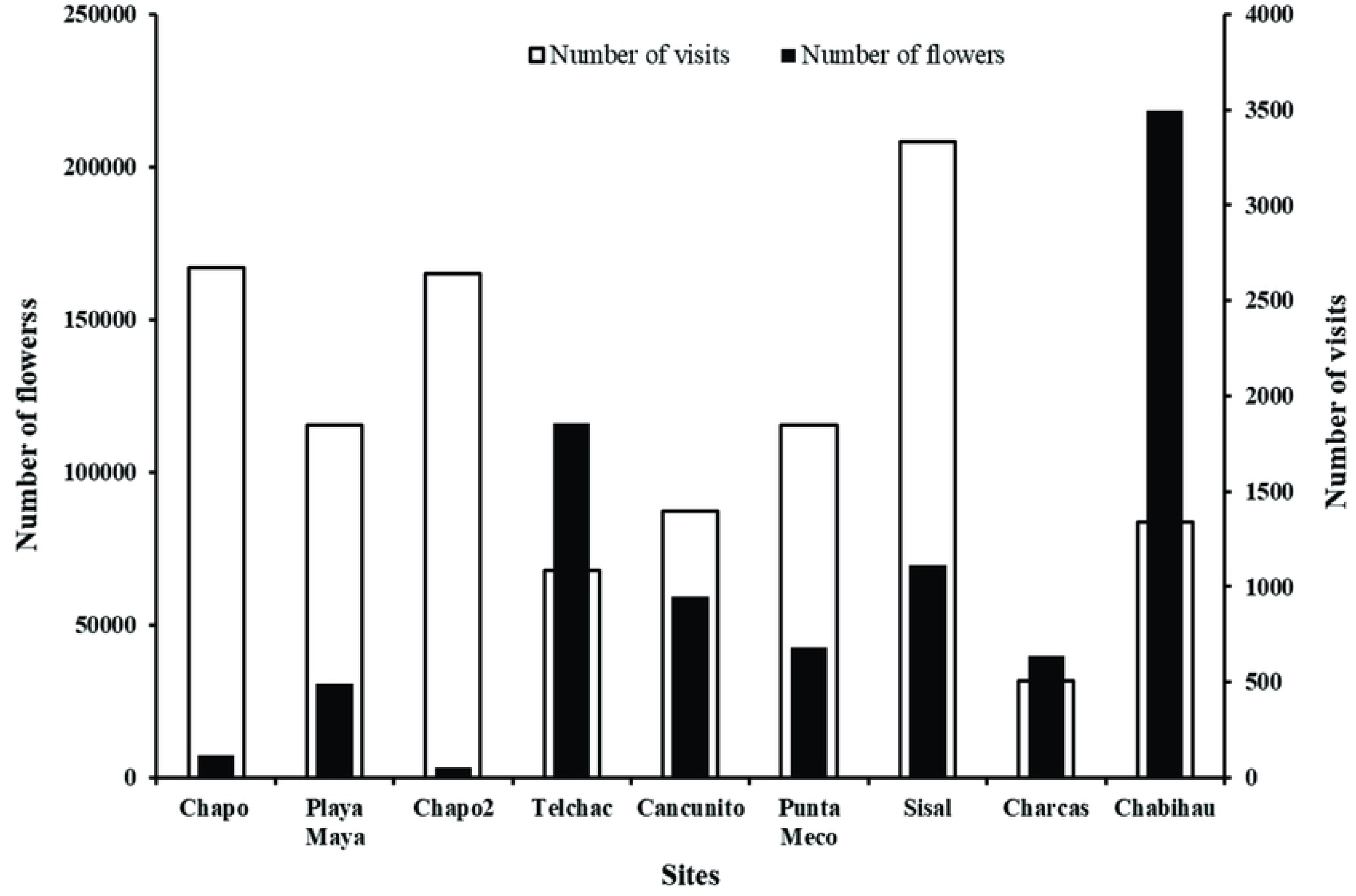
Number of total flowers and total visits registered in each study site. Sites are ordered according to increasing proportion of alien flower abundance (see T1).

The average percentage of alien flowers was high (75.7 ± 30.32), although we observed high among-site variation in the proportion of alien flowers (Table 1; range from 11% to 99%). A total of 14255 floral visits were recorded. The lowest number of total flower visits was observed at Charcas (504) and the highest at Sisal (3335) (Fig 1). Mean pollinator richness per plant species was highly variable among sites, being the highest at Playa Maya and the lowest at Sisal (Fig 2a). However, no significant differences were observed among sites or between native and alien plants (F ≤ 2.05, p > 0.05 in both cases). Although we observed significant differences among sites in pollinator species composition (PERMANOVA, *pseudo*-F_8, 26_ = 1.83, p < 0.05), no differences were observed between native and alien plant species (PERMANOVA, *pseudo*-F_1,26_= 0.7, p > 0.05). Furthermore, we did not find significant differences in pollinator visitation rate among sites (Fig 2b), or between alien and native species (F ≤ 0.83, p > 0.4 in both cases). However, pollinator species richness was significantly affected by flower abundance, with increasing number of pollinator species with increasing floral abundance (β= 0.37 ± 0.06, t_65_ = 6.23, p < 0.001). We also observed a significant negative effect of flower abundance on visitation rate, suggesting that increases in floral abundance leads to a decrease in pollinator visitation rate to individual flowers (β=-1.6 ± 0.17, t_65_ = −9.03, p < 0.001).

**Fig. 2.**
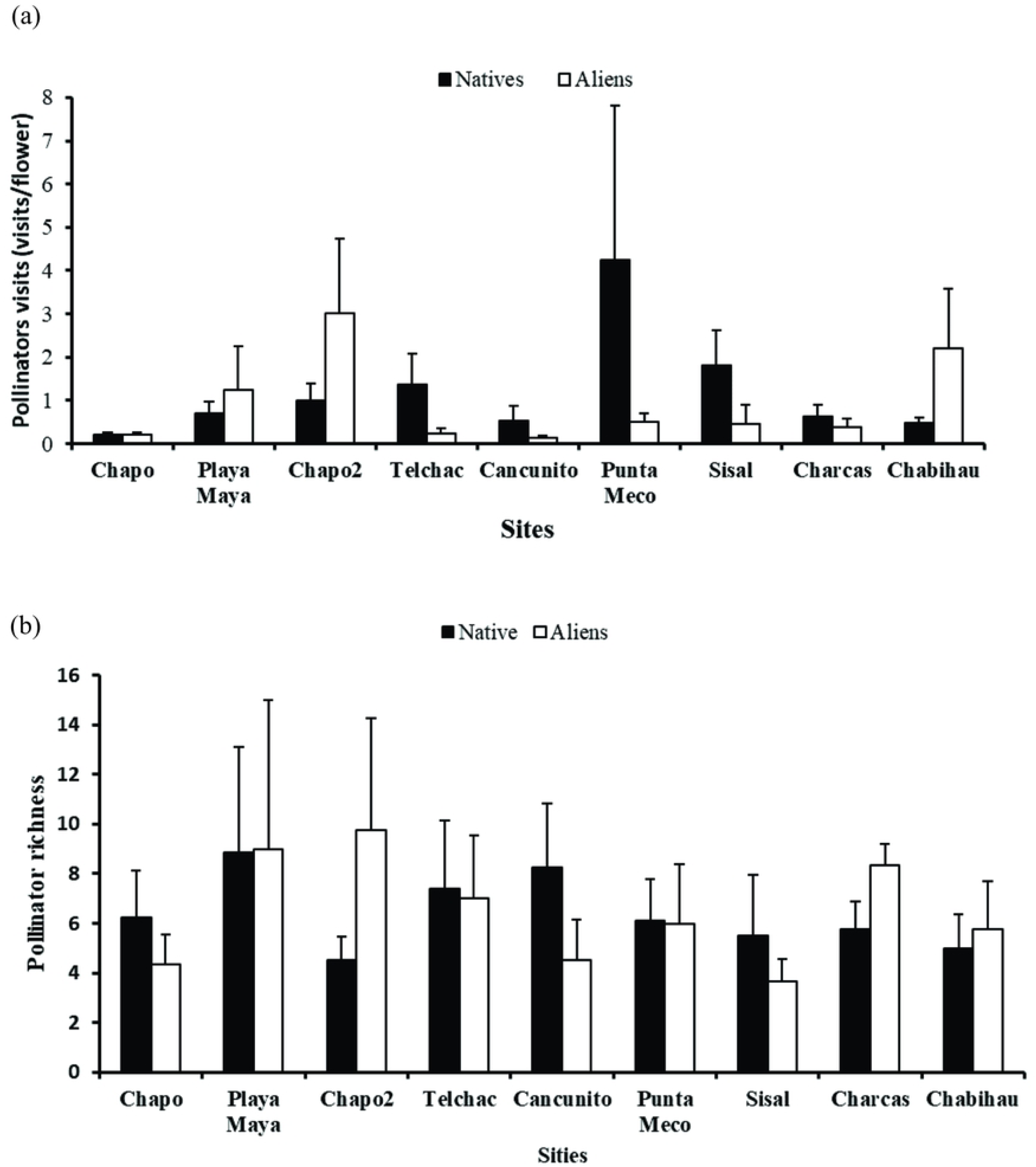
(a) Pollinator richness; and (b) flower visitation rate registered in each study site. Sites are ordered according to increasing proportion of alien flower abundance (see T1).

On average, across all communities, floral trait similarity was high (0.79 ± 0.1) and among-site variation was low (Table 1). Furthermore, we found no significant differences in floral traits across all sites between native and alien plants (F_1,33_ = 0.65, p = 0.42), nor within each community (F ≤ 3.7, p ≥ 0.07).

### Plant-pollinator networks

We recorded a total of 30 insect-pollinated plant species belonging to 19 families (S2 Table). A total of 73 insect species were recorded belonging to three orders: Diptera (26 species), Hymenoptera (27 species) and Lepidoptera (20 species) (S2 Table). Plant-pollinator interaction networks vary in size and contained between eight and 17 plant species and 22 and 38 pollinator species (Table 2; Fig 3).

**Table 2.** Number of plants and pollinator species and plant-pollinator interaction network metrics at nine sites across the north coast of the Yucatan Peninsula. Significant values of nestedness and modularity are shown in bold (P<0.05). Sites are ordered according to proportion of alien flower abundance (see Table 1).

| Sites | Number of plant species | Number of pollinators | Overall specialization network | Nestedness | Modularity | Robustness |
| --- | --- | --- | --- | --- | --- | --- |
| Chapo 1 | 12 | 31 | 0.53 | **25.80** | 0.43 | 0.64 |
| Playa Maya | 8 | 35 | 0.37 | **33.92** | 0.34 | 0.62 |
| Chapo 2 | 14 | 38 | 0.61 | **22.78** | 0.42 | 0.64 |
| Telchac | 14 | 36 | 0.35 | **27.81** | 0.35 | 0.68 |
| Cancunito | 8 | 22 | 0.52 | **36.66** | 0.36 | 0.66 |
| Punta Meco | 14 | 32 | 0.40 | **26.54** | 0.37 | 0.66 |
| Sisal | 11 | 27 | 0.29 | **26.75** | 0.38 | 0.62 |
| Charcas | 11 | 30 | 0.65 | 26.13 | 0.41 | 0.66 |
| Chabiahua | 17 | 28 | 0.37 | **27.8** | 0.36 | 0.69 |

**Fig. 3.**
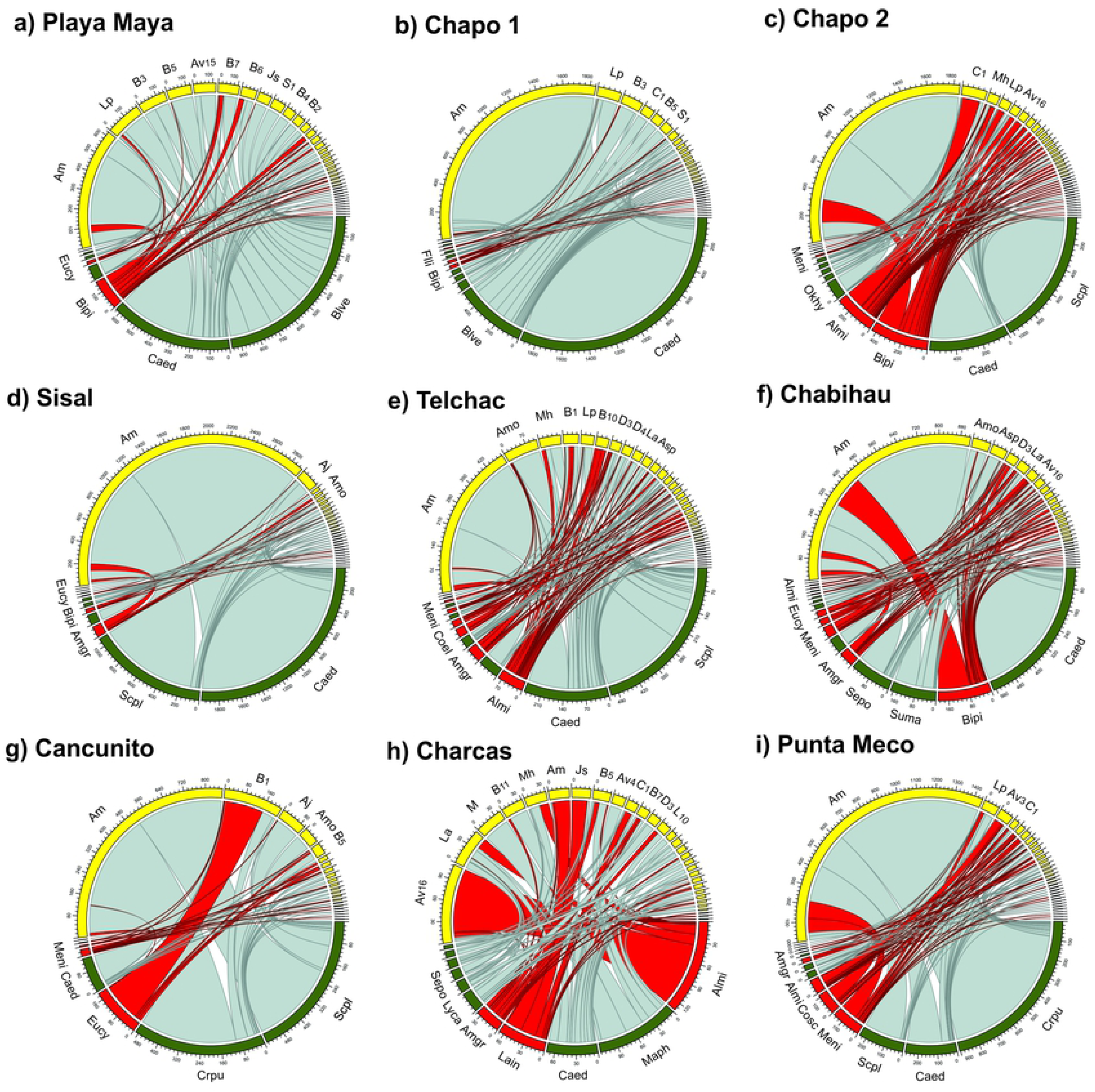
Plant-pollinator networks in each of nine sites (a-i) along the north of the Yucatan Peninsula, Mexico. Nodes in green and red represent native and alien plant species respectively. Nodes in yellow represent pollinator species (see TS1 and TS2 for a complete list of plant and pollinators and their codes). Numbers represent a scale for the number of interactions for a particular plant or pollinator species. Only codes for pollinators with more than 100 visits are shown. Red lines represent interactions between alien plant species and pollinators. Sites are ordered in increasing proportion of alien flower abundance (see T1).

However, in all sites, the highest percentage of visits corresponded to *Apis mellifera* (57.5%, range: 5.7% - 87.2%; with the exception of Charcas site, Fig 3). Overall, the most visited native plants were *Cakile edentula* and *Scaevola plumieri* (Fig 3) and the most visited invasive plants were *Bidens pilosa* and *Alternanthera microcephala* (Fig 3). Overall specialization (H2) (0.45 ± 0.12; range 0.29-0.65), nestedness (28.24 ± 4.31; range 22.78-33.66) and robustness (0.62 ± 0.02; range 0.62-0.68), varied little among sites (Table 2). All networks showed significant nestedness, (with the exception of one site) and non-significant modularity (Table 2). Other network metrics such as number of links per species, number of pairwise interactions and connectance are shown in S3 Table.

We did not find a significant relationship between alien species richness or the proportion of alien flowers and any of the observed structural network parameters (r ≤ 0.4, p ≥ 0.15 in all cases) suggesting that the ‘degree of invasiveness’ did not affect plant-pollinator network structure.

Species-level analyses showed that neither site nor plant origin (native or alien) affected species normalized degree and nestedness contribution (Table 3). In contrast, we observed a significant effect of plant origin on interaction strength (Table 3). However, this difference occurred only at one site (Chapo 2), where alien plants showed higher interaction strength than natives (t_47_ = 3.32, p = 0.002). Species-level strength and nestedness contribution significantly increased with increasing floral abundance (Table 3).

**Table 3.**
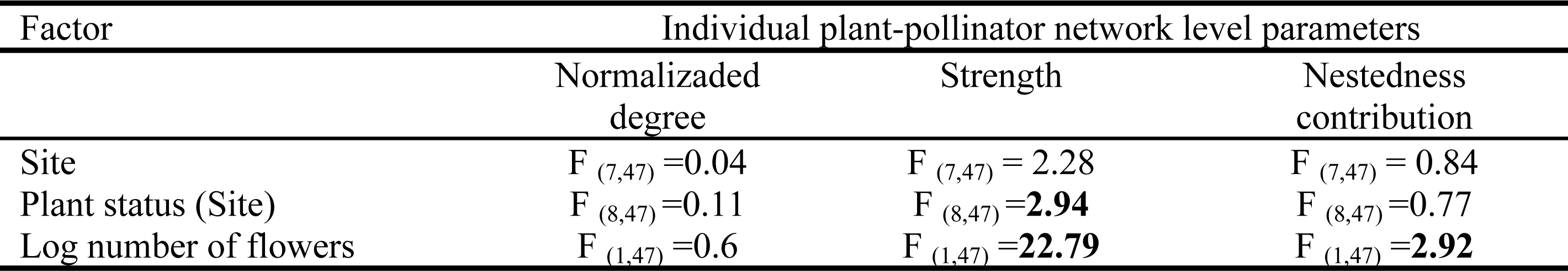
Results of mixed models evaluating differences in species-level plant-pollinator network parameters. Floral abundance (log number of flowers) was included as a covariate. Significant effects are shown in bold (p ≤0.01).

| Factor | Individual plant-pollinator network level parameters |  |  |
| --- | --- | --- | --- |
|  | Normalized degree | Strength | Nestedness contribution |
| Site | $F_{(7,47)} = 0.04$ | $F_{(7,47)} = 2.28$ | $F_{(7,47)} = 0.84$ |
| Plant status (Site) | $F_{(8,47)} = 0.11$ | $F_{(8,47)} = \mathbf{2.94}$ | $F_{(8,47)} = 0.77$ |
| Log number of flowers | $F_{(1,47)} = 0.6$ | $F_{(1,47)} = \mathbf{22.79}$ | $F_{(1,47)} = \mathbf{2.92}$ |

### Effect of alien species on plant-pollinator network structure via simulation of extinction scenarios

We observed significant differences in network specialization and nestedness among the different extinction scenarios (F_3,24 ≥_ 3.69, p < 0.05; Fig 4), but this was not the case for modularity (F_3,24_ = 0.55, p > 0.05). Overall network specialization in the ‘natives removed’ scenario was significantly higher compared to all other extinction scenarios and to the ‘intact’ network (t ≥ 3.5, p < 0.05, in all cases, Fig 4a) suggesting that the loss of native species may increase network specialization. The effect size of network specialization showed an 18% increase when native plants were removed compared to ‘intact’ communities (Table 4; S2 Figure).

**Table 4.** Mean (± SE) plant-pollinator interaction network metrics and average rate of change (Δ %, according to Hedge’s size effects) under different extinction scenarios in nine sites along the north coast of Yucatan, Mexico. Significant differences in the Δ % between “intact network” vs “aliens removed”; “intact networks” vs. “native removed”; and “intact networks” vs. “ransom removal” scenarios are shown in bold (p<0.05).

| Network topology estimator | Random models |  |  |  |  |  |  |
| --- | --- | --- | --- | --- | --- | --- | --- |
| | Intact networks | Aliens removed | ($\Delta$%) | Natives removed | ($\Delta$%) | Mixed removal | ($\Delta$%) |
| Overall specialization | 0.44 (0.05) | 0.41 (0.04) | -9.31 | 0.52 (0.06) | **+18.1** | 0.45 (0.05) | -1.0 |
| Nestedness | 28.24 (3.13) | 32.64 (3.62) | **+13.4** | 30.24 (3.36) | **+6.61** | 33.22 (3.69) | **+14** |
| Modularity | 0.38 (0.042) | 0.39 (0.043) | +2.6 | 0.39 (0.043) | +3 | 0.37 (0.041) | -1.0 |
| Robustness | 0.65 (0.007) | 0.62 (0.008) | **-4.8** | 0.63 (0.006) | **-3.8** | 0.62 (0.01) | **-4.8** |

**Fig. 4.** Structural plant-pollinator network parameters. (a) Overall network specialization (H2), (b) nestedness and (c) robustness for the observed plant-pollinator network (i.e. intact network) and the three extinction scenarios: “aliens removed”, “natives removed” and “random removal” (randomly exclusion of alien and natives).

We observed a significant decrease in network nestedness in the ‘intact’ network compared to the other extinction scenarios (t ≥ 3.1, p > 0.05 in all cases, Fig 4b), with the exception of the ‘natives removed’ scenario (Fig 4b). The effect size for nestedness showed a decrease of 19% in the ‘intact’ network compared to all extinction scenarios (Table 4; S2 Figure). However, the decrease in nestedness in the ‘aliens removed’ scenario was almost twice as high as in the ‘native removed’ scenario (13% and 6% respectively), suggesting that alien species have a greater effect on nestedness than native species (Table 4; S2 Figure). Comparisons between the ‘intact’ network vs. ‘native removal’ and ‘random removal’ scenarios showed that removal of plant species, regardless of their origin, significantly reduces network robustness (Table 4; Fig 4c). The effect size for modularity did not change significantly between any scenario (Table 4; S2 Figure).

### Effects of alien species at species level network

We found a significant effect of plant origin (nested in site) on normalized degree and strength (F ≥ 2.05, p<0.01 in both cases), but not for nestedness contribution (F_17,190_ = 1.45, p=0.11). However, for normalized degree, the results showed that any within-site comparison between native vs. alien species was significantly different (t ≤ 1.73, p ≥ 0.08, in all cases), and for strength the results only showed significant differences between native vs. alien species at one site (Chapo site; t_270_= 3.14, p<0.01). In contrast, we found a significant scenario effect on normalized degree, strength and nestedness contribution (F ≥ 2.05, p<0.01 in all cases). Interestingly, for normalized degree and strength we found significant differences between the ‘intact’ network and the ‘native removal’ and ‘alien removal’ (t_270_ ≥ −2.65, p < 0.01 in both cases), suggesting species removal diminished normalized degree and strength regardless of plant origin (native or alien). For nestedness contribution we found similar results, the ‘intact’ network showed significant differences with ‘alien removal’ (t_190_= 2.82, p < 0.01), and marginally significant differences with ‘native removal’ scenarios (t_190_= 1.9, p = 0.059). Furthermore, for none of the species-level parameters we found significant differences between native or alien ‘removal’ scenarios (t ≤ 0.7, p ≥ 0.4 in all cases).

## Discussion

Our results show that none of the evaluated network parameters is affected by increasing ‘intensity’ of plant invasion (alien species richness and flower abundance), suggesting that even low alien species richness or flower abundance can have significant impacts on native plant and pollinator communities. Our results also suggest that alien species are well integrated in native plant-pollinator networks (Fig 3), and that pollinator use overlap with natives is high, which is likely mediated by high levels of floral trait similarity between alien and native species. Consistent with these results our simulated extinction scenarios suggest that alien species play an equivalent role to natives in network structure and stability in our studied coastal plant communities. These and other results are discussed in detail below.

Contrary to our expectations, and in spite of extensive among-site variation in alien species richness and floral availability, we did not observe among-site differences in any of the network structural parameters suggesting that the plant-pollinator network structure is not affected by the ‘intensity’ of species invasion. It is possible that this lack of effect may be mediated by the high abundance of the introduced honey bee *A. mellifera* at our study sites, which contributes disproportionally to flower visitation at all sites (ca. 60% of visits; see Fig 3). It has been shown that ‘super generalist’ pollinators such as honeybees can facilitate the integration of alien plants into native pollination networks and support the structure of the network in the presence of alien species [11, 16]. Thus, the high incidence of *A. mellifera* at our study sites may be a key factor mediating plant-pollinator network structure regardless of alien species richness and flower abundance at a site (i.e. intensity of invasion). Interestingly, in our study, the only community that did not show a significant nested structure (Charcas site; Fig 3) was also the one with the lowest proportion of *A. mellifera* visits (5.7%), lending support to the prediction that this ‘super generalist’ pollinator plays an important role in network structure. It is important to note that low statistical power (nine communities) could also had hampered our ability to observed significant effects (Table 1) and thus studies that evaluate the effect of ‘invasion intensity’ on network structure over a wider range of communities and ‘intensities’ of invasion are needed. Nonetheless, our results suggest that invasive species are well integrated within native plant-pollinator networks and their effect on network structure could be considered ‘equivalent’ to that of native plant species. There are three lines of evidence that support the ‘equivalency’ in the functional role of native an alien plants species within plant-pollinator networks in our study system: 1) comparisons of our simulated extinction scenarios suggest that the loss of alien plant species has the same effect on network structure and robustness as the loss of native species. 2) No differences were observed in any of the species-level network parameters (i.e., normalized degree, strength and nestedness contribution) between alien and native plant species and 3) floral trait similarity between native and alien was notably high (≥ 77%). The capability of alien plant species to fully integrate into native plant-pollinator networks has also been observed in other ecosystems [5, 10, 17].

Interestingly, simulation analyses revealed a significant difference in network specialization between the ‘natives removed’ and all the other extinction scenarios. This suggests that the loss of an equivalent number of native species (relative to alien species) would result in an increase in network specialization. This result may suggest that alien species establish relatively ‘specialized’ interactions with pollinators already present at a site [11, 17, 59]. For instance, in some sites alien plants were visited only by one or few insect species (see Fig 3). *Euphorbia cyathophora* was visited only by one (*Crysntrax dispar;* Bombyliidae) and two pollinator species (*Bombilide spp* and *Syrphidae spp*) at Chapo1 and Sisal sites respectively. *Amaranthus greggii* was only visited by *Junonia scenia* (Nymphalidae) and *Apis mellifera* at the Canunito site. This is contrary to the expectation that generalized pollination systems are favored in alien species in order to be successful. Thus, a more detailed analysis of the degree of pollinator specialization and generalization in native and invasive plant species and pollinators in our communities is underway. Furthermore, the observed reduction in network nestedness in the three scenarios in which aliens, natives or random species were removed compared to ‘intact’ networks suggests that an overall loss of species will decrease network robustness and increase species vulnerability to extinction [13, 14, 57]. In fact, the effect size analysis showed that removal of plants species, regardless of their origin (native or alien) significantly reduced network robustness. Empirical studies have shown similar results (reviewed by Stout and Tiedeken [12]. For instance, in a study in multiple invaded and non-invaded communities across Europe, Vilà et al. [60] found that the plant-pollinator networks appear to be permeable and robust to the introduction of invasive alien species. Padrón et al. [21] found that the alien genus *Opuntia* did not affect plant-pollinator network nestedness and connectance in the Canary Islands and the Balearic Islands. These results suggest that even though the arrival of alien species into the studied coastal plant communities has dramatically increase in the past 30 years [32], these seem to be well integrated and significantly contribute to maintain plant-pollinator network structure and robustness.

On the other hand, we did not find evidence of invasive species effects on species-level parameters despite previous evidence from other systems suggesting the contrary [21, 57, 59]. For instance, it could be expected that if alien species have a generalized pollination system they may contribute more to increased nestedness. It has been shown that species that contribute more to nestedness are also more important for the persistence of the entire network [55]. However, our results suggest that alien species are also capable of establishing specialized interactions with pollinators (see above) and thus natives and alien’s plants contribute equally to nestedness. Species-level strength and nestedness contribution, however, were positively affected by overall flower abundance suggesting that resource availability mediates the diversity and strength of plant-pollinator interactions in the studied coastal communities regardless if these are from native or invasive plant species [61].

The integration of alien species into the studied plant-pollinator communities could be facilitated, at least partially, by high floral trait similarity with natives, which allows the use of existent pollinators in a community [24]. It has been proposed that floral trait similarity between native and alien species could mediate the effect alien species on the pollination success of natives [7]. For instance, some studies have found that alien plants that share some floral traits with native plants inflict stronger negative pollinator-mediated impacts on natives compared to invasive species that do not share similar floral traits [15, 24]. This suggests that floral traits of co-flowering species may more strongly underlie effects on pollination success of co-flowering neighbors rather than plant origin (i.e., native or alien; [41]). The high floral similarity may also help explain the ‘equivalence’ in the functional role of native and alien plants within the studied plant-pollinator networks (see discussion above). However, it must be pointed out, that high floral similarity can also lead to high flower visitor overlap between native and invasive species resulting in heterospecific pollen transfer [15, 62]. Thus, future studies should also consider the potential effects of alien plant species on the ‘quality’ of the interaction between native plants and their pollinators (e.g. heterospecific pollen transfer networks; [63, 64]. Preliminary data of pollen transfer between native and invasive plants in our study showed that alien heteroespecific pollen represents more than 10% of total pollen load on native stigmas (Parra-Tabla unpublished data). Thus, integration of the alien plants into native networks could have indirect detrimental effects on fruit and seed production in native plants [65]. Hence, we emphasize the importance of evaluating invasive species effects at multiple levels of the pollination process [7, 12, 66, 67] in order to advance our understating of their effects in natural communities.

Finally, network nestedness values observed here were lower than those typically reported for other plant-pollinator networks [46, 52] which may be the results of a relatively small network size at our study sites (<50 species; [51]. Yet, even though the number of interacting species in our communities was relatively low (min 38 and max 52 species), nestedness was statistically significant in eight out of nine sites. It is possible that in these ecosystems, which can be considered ‘stressful pollination environments’ (i.e., strong winds and high temperatures; [40, 41, 59], plant-pollinator network structure is less dependent on size because of a stronger interdependence between plant and pollinators, which allows them to persist in these harsh environments [22, 59]. For instance, average visitation rate to flowers was very low across all coastal co-flowering communities (ca. 1 visit/flower/min; min 0.32 - max 2.64) and pollinator visitation rate further decrease with increasing overall flower abundance, thus suggesting that plant reproduction in these coastal communities may be limited by pollinator availability [40]. Similar results have been observed in other coastal [40] and insular invaded communities [16] with few participating species. Thus, our results emphasize the importance of considering the importance of not only the number, but the strength and type of plant-pollinator interactions established within communities, in mediating network structure.

## Acknowledgements

We thank Kevin Baas, Rigel Silveira, Brian Suárez, and Daniel Tzuc for their valuable help in the field-work.

## Supplementary material

S1 Table Plant species list

S2 Insect species list

S1 Figure. Location of the nine study sites along the north of the Yucatan Peninsula, Mexico.

S2 Figure. Size effects (CI 95%) for overall specialization, nestedness, modularity and robustness contrasting different scenarios with intact networks, (a) “Aliens removed”, (b) “natives removed”, and (c) “mixed removal”.

Data set Excel file.

